# Nightmares share strong genetic risk with sleep and psychiatric disorders

**DOI:** 10.1101/836452

**Authors:** Hanna M Ollila, Nasa Sinnott-Armstrong, Katri Kantojärvi, Teemu Palviainen, Anita Pandit, Robin Rong, Kati Kristiansson, Nils Sandman, Katja Valli, Christer Hublin, Jaakko Kaprio, Richa Saxena, Tiina Paunio

## Abstract

Nightmares are vivid, extended and extremely dysphoric dreams that awaken the dreamer. Twin studies indicate that nightmare frequency has a heritability between 36 and 51% providing evidence for genetic factors underlying predisposition to nightmares. Furthermore, while cross-sectional epidemiological findings suggest that heavy alcohol usage, traumatic experiences and psychiatric diseases associate with nightmares, the causal relationships between these conditions and nightmares have remained unknown. To examine the biological mechanisms behind nightmares, we performed a genome-wide association study in 28,596 individuals from Finland and the United States. We identified individual genetic variants that predispose to nightmares near *MYOF* (rs701873, p=2.18e-8) and *PTPRJ* (rs11039471,p=3.7e-8), a gene previously associated with sleep duration. We found a strong genetic correlation between the frequency of nightmares and traits related to personality and psychiatric disorders; neuroticism (rg=0.59, p=8e-7), post-traumatic stress disorder (PTSD) (rg=0.58, p=0.004) as well as major depressive disorder (rg=0.68, p=7e-4), and sleep traits; and daytime sleepiness (rg=0.62, p=1e-6) and insomnia (rg=0.50, p=1.87e-5). Analysis of directionality using mendelian randomization showed a significant effect from feelings of fed-up (p=0.001), nervous (p=0.004) and miserableness (p=0.0045) to nightmares with no evidence of pleiotropy and no evidence of nightmares predisposing to psychiatric or psychological problems. Our findings suggest that nightmares are caused by unique genetic risk factors, and here we identify the first individual genetic associations. In addition, a substantial effect on nightmares is conveyed through underlying psychological and sleep problems, with psychological problems being causal for nightmares.

## INTRODUCTION

Nightmares are vivid, extended and extremely dysphoric dreams that usually include themes involving threats to survival, security, physical integrity or self-esteem, and cause the dreamer to wake up (American Academy of Sleep Medicine). While sporadic nightmares and bad dreams are common and generally harmless, a subset of nightmares reflects underlying pathology. Indeed, while almost everyone experiences sporadic nightmares, experiencing regular nightmares is relatively rare, with less than 5% of the overall population and with higher prevalence in females compared to males, as well as a higher prevalence of nightmares in childhood compared to adulthood ^1-3^.

While the risk factors behind nightmares are not well understood, nightmares can be caused by negative experiences and fear such as traumatic events, as seen in war veterans with PTSD ^2^. We have previously studied the epidemiological correlates of nightmares extensively. These and other epidemiological studies have shown that both sleep problems and severe sleep disorders, such as insomnia and narcolepsy, are associated with nightmares, there being a strong association with insomnia, fatigue and sleep terrors ^4-7^. Furthermore, we have previously shown a familial aggregation between nightmares and psychiatric disorders ^3^. Similarly, we and others have previously shown that nightmares correlate with sleep and psychiatric traits, alcoholism, insomnia and overall pain, including headache ^2,3,8,9^. However, the biological basis behind nightmares or the causality between nightmares and comorbid traits are not known.

Interindividual genetic differences account for 36 to 51% of the variance in liability to nightmares in the population ^3^. Genetics offers an opportunity to identify new predisposing loci, understand whether underlying biological mechanisms are shared with psychiatric or other traits, and identify which associated factors contribute causally to frequent nightmares. Indeed, all earlier studies sketch a pathway where nightmares overlap with psychiatric and sleep traits. However, whether they are a cause or consequence of nightmares has remained inconclusive; potentially other confounders including shared genetic factors may account for the observed association.

Previously, in a longitudinal study of war veterans with PTSD it was seen that both insomnia and nightmares had a strong association with PTSD but insomnia did not resolve by itself whereas nightmare symptoms got milder. This suggests a stronger independent component from insomnia for PTSD symptoms than nightmares ^10^. Furthermore, nightmares tend to remain unreported in the clinical setting. Similarly, we also know that people underestimate the frequency of nightmares in retrospective analysis versus daily logs ^11^. Consequently, the overall clinical relevance of nightmares has probably been underestimated despite their known epidemiological association with sleep disorders, psychiatric disorders, and suicide ^12^.

In this study our goal was to examine the underlying biological and epidemiological mechanisms that affect the frequency of nightmares. We aimed to understand the putative causal links between nightmares, sleep traits and psychiatric traits. Given that nightmares are heritable, we used a genome-wide association study design (GWAS) in three cohorts with self-reported frequency of nightmares to elucidate underlying biology, shared mechanisms and links with psychiatric diseases.

## METHODS

### Study populations

28,596 individuals participated in this study from Finrisk (N=21,223), Finnish Twin Cohort Study (N=5,556) and Genes for Good (N=1,797). The individuals contributed genetic and questionnaire data and provided a written informed consent for research.

#### FINRISK (n=21,223)

This cohort comprises health surveys, collected every five years, from cross-sectional random population samples of Finnish adults. The data is collected in forms of health questionnaires and a formal health examination at a local health care center. In the Finrisk dataset, nightmares were assessed with a question “During the past 30 days have you had nightmares?”. The response options were “often”, “sometimes” and “never.” Other measures, including age, gender and the survey year that the subject took part in (Finrisk surveys 1997, 2002 or 2007) as well as alcohol use, were assessed. Alcohol intake was measured using a question on frequency of alcohol intoxication. Nightmares were used as a linear trait from 1 to 3 with 1 being no nightmares and 3 frequent nightmares. Genotyping and genome-wide imputation was done as part of the Finrisk data analysis using Sequencing Initiative Suomi (SISu) project data as reference. Genome-wide association analysis was conducted using SNPtest with option *score* in the SNPtest package ^13^. Analyses were adjusted for relevant covariates comprising age, gender and identity by descent calculated genetic correlation matrix with top 10 principal components, the survey year, and genotyping chip. In addition, individuals that were intoxicated once or more often per month were removed in the secondary analysis, as frequent heavy use of alcohol has been previously found to increase risk for frequent nightmares ^4^.

#### Finnish Twin Cohort (n=5,556)

The study cohort consists of same-sexed twin pairs born before 1958, who participated in two questionnaire surveys in 1975 and 1981. In 1990, twins who had participated in either previous survey and who were born between 1930 and 1957 were invited to participate in a questionnaire survey ^14^. The 1990 survey included a broad set of items on frequency of parasomnias in childhood and adulthood, as reported earlier ^3,15^. In the Finnish Twin Cohort, two questions assessed the frequency of nightmares in childhood and adulthood, respectively ^7^. In the analysis, we used the data on nightmares as adults. For adults, the question was “How often have the following nighttime symptoms been present in adulthood Alternatives were ‘weekly’, about once a month, less often, never and cannot say. Those with weekly nightmares were defined as cases and those with nightmares never, rarely or monthly as controls. Heavy drinking occasions were defined as consuming ‘at least once a month and on a single occasion’, more than five beers, a bottle of wine or a half-bottle of spirits (or a similar amount), to correspond to five standard drinks or >60 g of pure alcohol on a single occasion ^16^. For Finnish Twin Cohort we used Haplotype Reference Consortium release 1.1 reference panel ^17^ for genotype imputation using protocol provided by University of Michigan Imputation Server ^18^. For Finnish Twin Cohort: Primary analyses, analyses were conducted using RVTESTS ^19^ with linear mixed model regression using score test for testing association. The sample relatedness and population stratification has been taken into account in genetic relatedness matrix used as a random effect of the model.

#### Genes for Good (n=1,797)

Genes for Good is an online genetics study conducted through a Facebook web app. Participants self-report health and behavioral data through online surveys and submit a saliva sample in the mail for genotyping. The sample includes participants over 18 years of age from all 50 U.S. states. Altogether, 1797 individuals in Genes for Good participated in the study. In Genes for Good the frequency of nightmares were assessed with a question: “During the last 30 days, how often have you had nightmares? Nightmares are dreams that evoke so strong negative feelings that they wake you up.” Answering options were: “1. Almost every morning, 2. Several times a week, 3. Once a week, 4. Two or three times a month, 5. Once a month, 6. I did not have nightmares that woke me up during the last month”. Phenotypes were harmonized similar to Finnish Twin cohort: Those with nightmares at least once a week were defined as cases and those with nightmares never, rarely or less than weekly as controls. Analysis was computed as or logistic mixed model regression with SAIGE ^20^, adjusting for age, sex and principal components. In the secondary analysis, as information was not available on how much alcohol individuals consumed at one occasion, use of alcohol was adjusted instead as many drinks people have had during the last 30 days.

#### Meta-analysis

Fixed effect meta-analysis was used to combine effects across cohorts using META ^21^ and compared to those using METAL with sample size weighted analysis ^22^.

#### Biological consequences of genetic variants

Downstream analyses of gene level overlap with other traits and pathway analyses were calculated using MAGMA with variants under p-value threshold 1e-5 as implemented in FUMA, and single Cell Type enrichments were calculated with all available cell types in Mouse Cell Atlas using MAGMA as implemented in FUMA. http://fuma.ctglab.nl/ ^23^.

#### Genetic correlation

In order to estimate the genetic correlation between two traits, we used LD score regression, which considers linkage disequilibrium (LD) (e.g. correlation due to close proximity at the DNA chain) between genetic variants. Similarly, trait heritability explained by common variants and tissue specific partitioned heritability were estimated using LD score regression ^24^. P-values were Bonferroni corrected.

#### Mendelian Randomization

Mendelian Randomization was computed between nightmares and those traits that showed significant genetic correlation with nightmares. Analysis was performed using MRCIEU/TwoSampleMR package in R 3.5.0. We calculated inverse variance weighted and MR-Egger models to estimate effect sizes. For those associations with significant effect we estimated pleiotropy as implemented in the MRinstruments R package. For nightmares with only two significant genome-wide SNPs a lower threshold of 1e-5 was used to extract exposure instruments clumping individual lead variants per locus together in order to adjust for LD.

## RESULTS

### Meta-analysis of GWAS of nightmares identifies genetic loci at PTPRJ and MYOF

We have earlier studied the epidemiological correlates of nightmares extensively in the cohorts examined here^2-4,7,8^. In the current study, we explored whether there were independent genetic risk factors for frequency of nightmares. Overall SNP based heritability was relatively modest [SNPh^2^=0.05 se=0.017]. In primary meta-analysis of GWAS in 28,596 subjects, we identified a significant association with nightmares for a locus intronic to *PTPRJ* (rs11039471, p=3.7e-8) (**Figure 1, Supplementary Figure 1-2, Supplementary Table 1**). Previously, an independent signal at the same locus, not driven by the same lead variants, associated with total sleep duration, short sleep duration and glycemic traits ^25,26^ suggesting a broader pleiotropic role for the *PTPRJ* locus in regulating sleep and symptoms of disturbed sleep such as nightmares.

**Figure 1.**
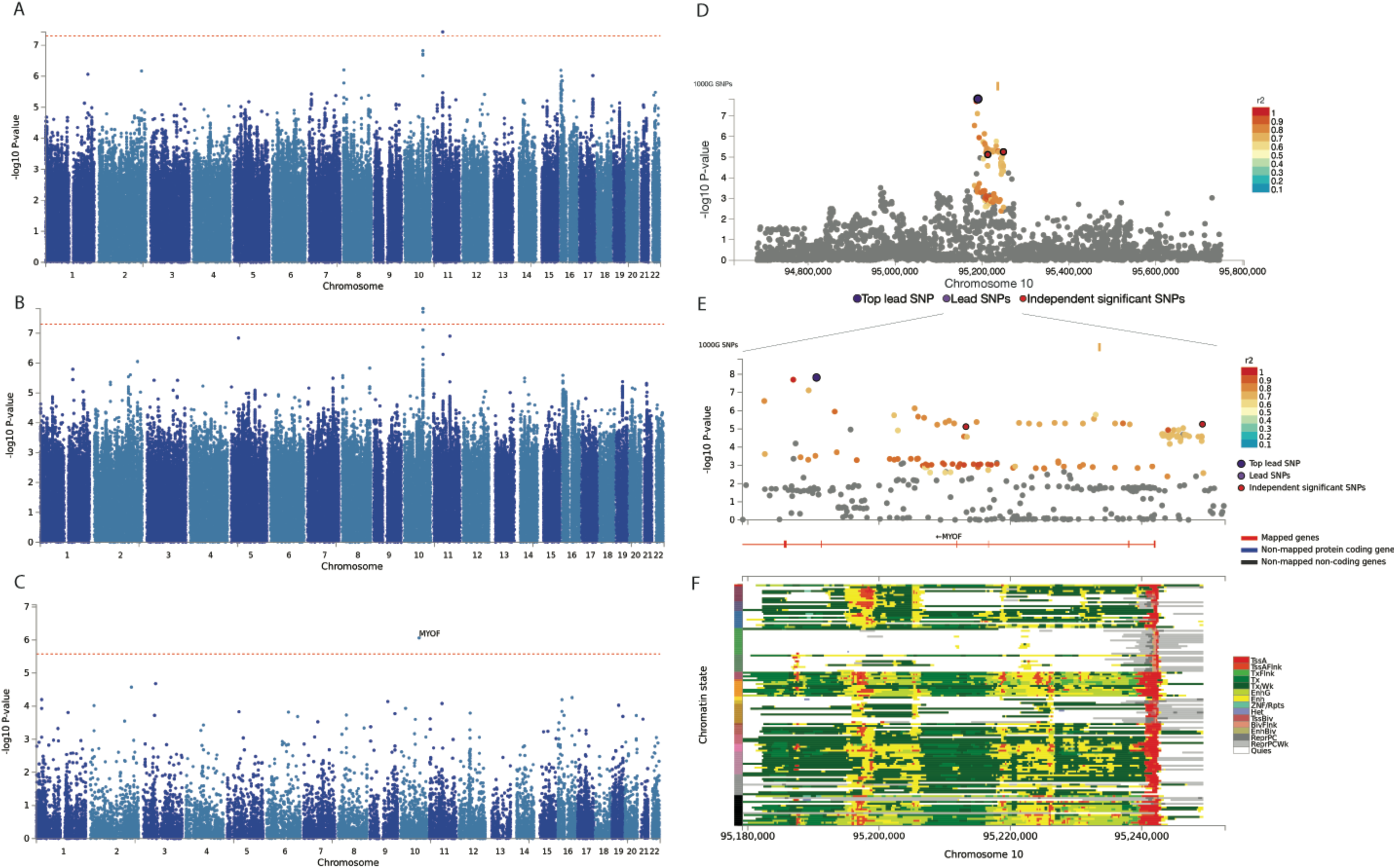
Single point association with nightmares. The genetic association with A) nightmare frequency in model adjusted for age, gender, population stratification and batch effects or B) nightmare frequency in model adjusted for age, gender, population stratification and batch effects excluding individuals who get intoxicated at least once per month shows significant association with *MYOF* locus. C) Gene based test for nightmare frequency shows significant association with *MYOF* locus. D) Regional association of *MYOF* locus and E) zoomed in association with *MYOF* locus show that E) the region is ubiquitously transcribed except for brain tissues (marked green on Y-axis).

Previous studies and our observations above show that heavy use of alcohol correlates with nightmares ^4,27^. Therefore, we explored the genetic determinants after removing individuals that were intoxicated at least once per month as a secondary analysis. This analysis identified an additional signal intronic to *MYOF* (rs701873, p=2.18e-8, Figure 1B) with two additional GWAS significant SNPs (rs701874 and rs788094) in tight LD. Multiethnic analysis with Genes for Good strengthened this association (p=1.2e-8, Supplementary Table 1). Furthermore, gene-based test supported association of *MYOF* (Figure 1C). The associating variants are intronic to *MYOF* - a gene normally expressed in microglia in the brain as well as in high levels in the bladder.

### Tissue enrichment analyses implicates brain and neuronal tissues in nightmares

To explore downstream biological mechanisms we computed tissue enrichment for closest genes to 29 lead variants with p-value <1e-5 using GTEx data. This analysis showed a global enrichment of genetic loci active in the brain likely reflecting the associations with sleep, mood and alcohol consumption as well as direct effects on nightmares themselves (**Figure 2**). Furthermore, we saw association with genes expressed in the bladder (**Supplementary Figure 2**) suggesting a separate role for mechanisms involving sleep interruption and potentially, getting up during the night to use the restroom. Similarly, single cell enrichment identified both brain cell types: Astrocytes (p-value = 6e-3) as well as Bladder cells enriched with nightmares (p=5e-4) (**Supplementary Table 2-3**).

**Figure 2.**
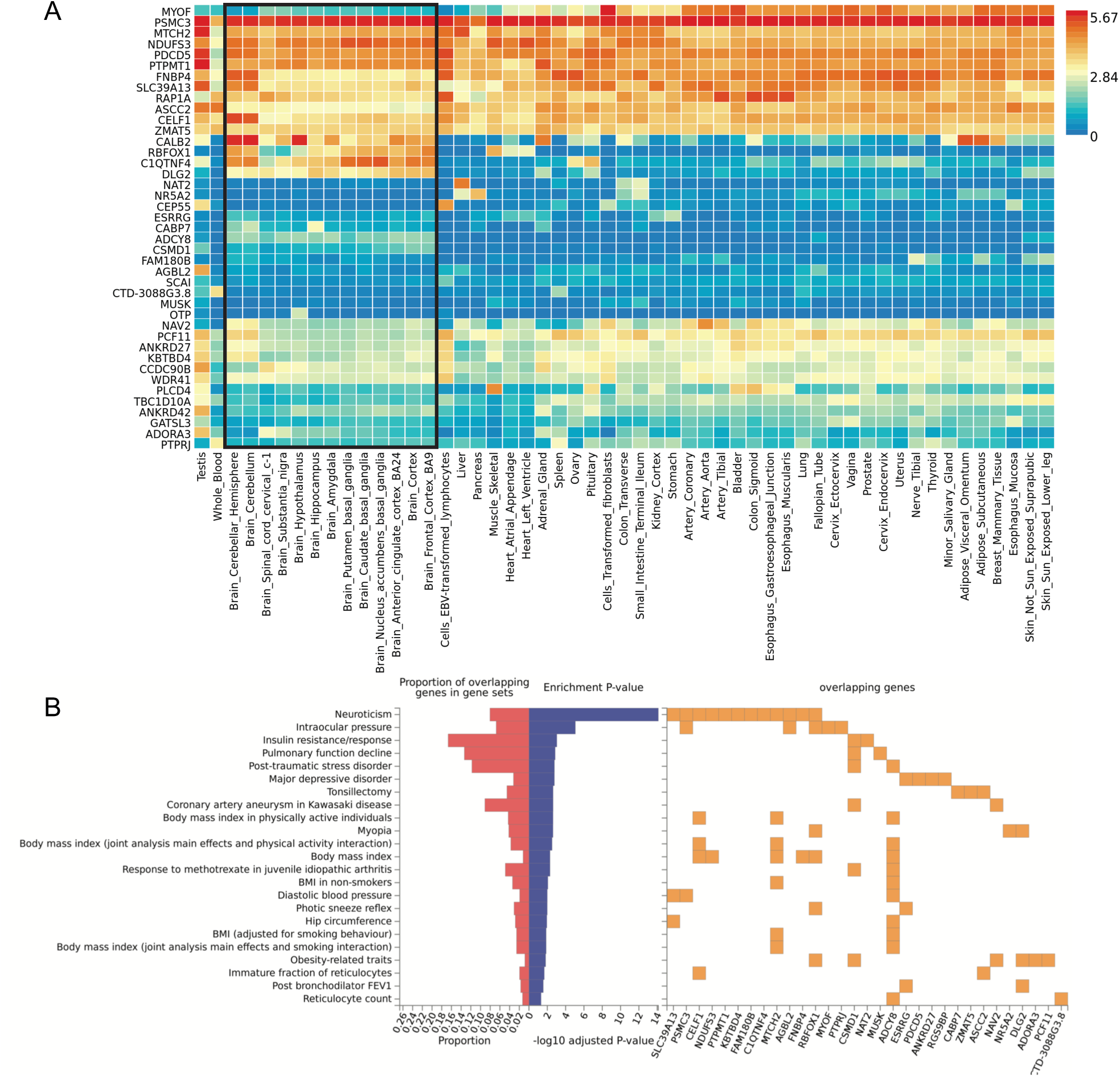
Expression of nightmare target regions and overlap with previously explored traits. A) Expression of genes in loci with p<1e-5 using GTEx v6. Brain tissues are highlighted and enriched for expression. B) Overlap of nightmare loci with p<1e-5 (n=29 loci) with variants in same genes from previous association studies.

### Nightmares overlap with neuroticism and psychiatric disorders at gene level

To explore whether the same genes contribute to nightmares and psychiatric traits, we examined the genetic loci (n=29 loci) that associated with frequency of nightmares with p<1e-5. Variants within these most significant loci overlapped significantly with genes at loci that are significant for other psychiatric traits or personality traits related to such problems (**Figure 2, Supplementary table 4**). Most notably, these traits include neuroticism (p corrected= 3.53e-18 with *SLC39A13, PSMC3, CELF1, NDUFS3, PTPMT1, KBTBD4, KBTBD4, FAM180B, C1QTNF4, MTCH2, AGBL2, FNBP4, RBFOX1*), PTSD (p adj=2.42e-3, with *CSDM1* and *ADCY8*), and major depressive disorder (p corrected < 0.05 *ESRRG, PDCD5, ANKRD27* and *RGS9BP*).

### Nightmares show genome-wide genetic correlation with sleep and psychiatric traits

We first examined the genetic correlation between PTSD^28^ and frequency of nightmares. We discovered a significant correlation (rg=0.58 p=0.004) evidencing for shared genetic vulnerability between frequent nightmares and PTSD. To assess whether previously suggested association between nightmares and psychiatric traits is mediated by genetic factors we computed genetic correlation between the GWAS of nightmares and publicly available summary results of GWAS for psychiatric disorders. We saw a strong genetic correlation of nightmares with major depression (rg=0.68, p=0.0007) and symptoms of depression (rg=0.58, p=0.0001), and an inverse association with subjective well-being (rg=-0.62, p=3e-5) (**Table 1).** Interestingly, pointwise association was also seen with schizophrenia (rg=0.25, p=0.015, p corrected = ns.) but not with bipolar disorder (rg=0.027, p=0.83). The correlation with schizophrenia did not sustain correction for multiple testing (**Supplementary Table 5**).

**Table 1.**
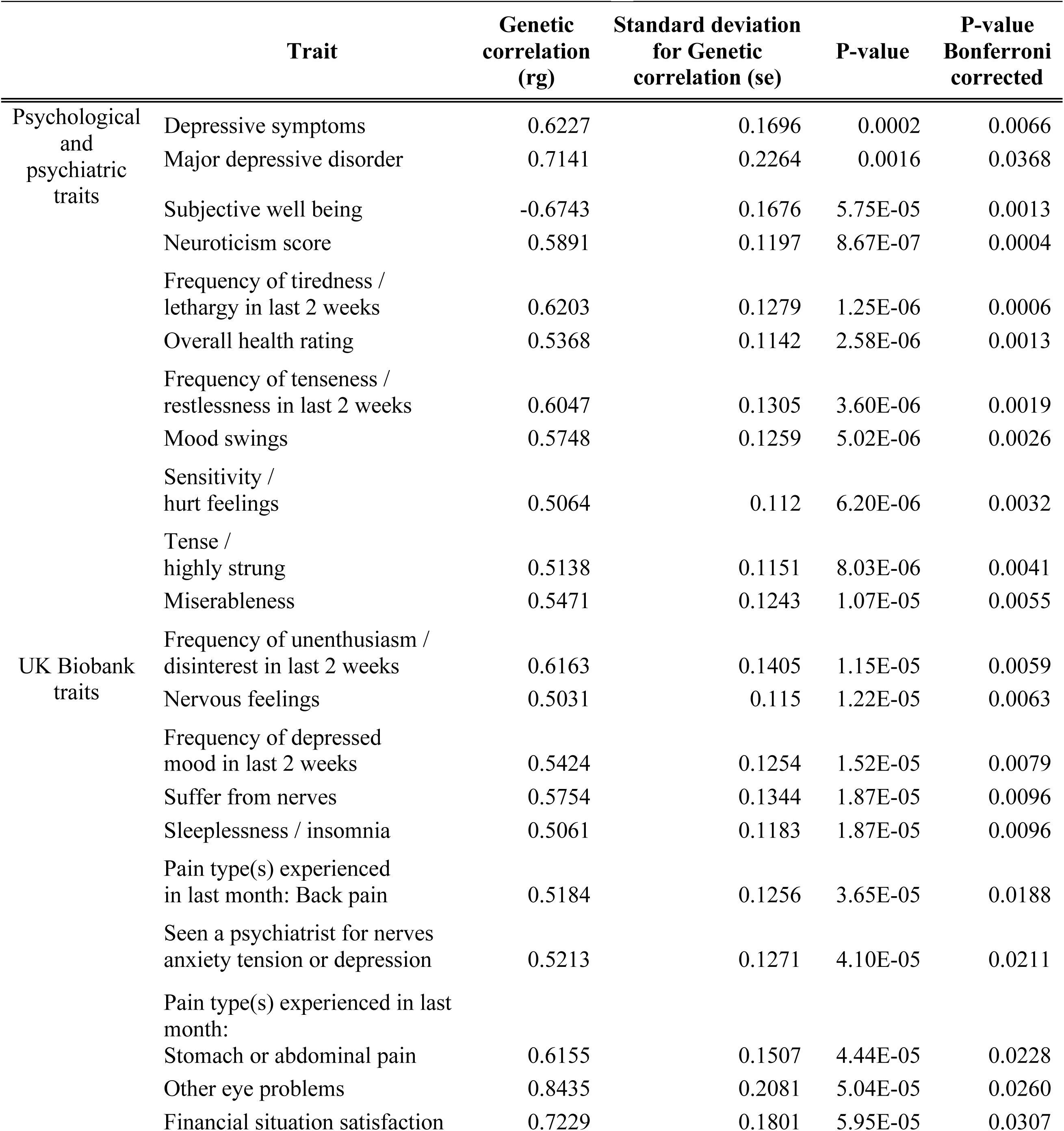
Genetic correlation from GWAS of Frequency of nightmares.

**Table 2.** Single nucleotide polymorphisms associating with the frequency of nightmares across cohorts.

| Gene,<br>MarkerName | Chr | bp | Ref<br>allele | Effect<br>Allele | Minor Allele<br>frequency | Finnish cohorts |  |  | Multiethnic |  |  |
| --- | --- | --- | --- | --- | --- | --- | --- | --- | --- | --- | --- |
|  |  |  |  |  |  | Effect | StdErr | p-value | Effect | StdErr | p-value |
| Age, sex and PC adjusted |  |  |  |  |  |  |  |  |  |  |  |
| PTPRJ, rs11039471 | 11 | 47975794 | C | G | 0.161 | 0.0413 | 0.0075 | 3.71E-08 | 0.0407 | 0.0075 | 5.37E-08 |
| Age, sex, alcohol and PC adjusted |  |  |  |  |  |  |  |  |  |  |  |
| PTPRJ, rs11039471 | 11 | 47975794 | C | G | 0.161 | 0.0407 | 0.0079 | 2.65E-07 | -0.0403 | 0.0079 | 3.28E-07 |
| MYOF, rs701873 | 10 | 95190568 | C | T | 0.3864 | 0.0331 | 0.0059 | 2.15E-08 | 0.0336 | 0.0059 | 1.24E-08 |

*Chromosome (chr) and Base pair (bp) positions in hg19. P-value calculated with standard error (StdErr), Effect. Fixed effects meta-analysis.*

In order to examine a broader set of human traits to find potentially stronger and novel correlates for nightmares, we then examined the genetic correlation of nightmares with 515 traits measured in UK Biobank, including sleep and psychiatric traits as well as additional autoimmune, lipid, cardiometabolic and anthropometric traits. The results were similar to the previously observed phenotypic association with sleep and psychiatric disorders with the strongest associations seen for psychiatric traits neuroticism and nervous feelings, frequency of tenseness, overall health rating, mood swings, sensitivity and lack of enthusiasm (p<1e-4 and rg>0.5 for all traits). In addition, there was a significant correlation with sleep traits: frequency of tiredness and symptoms of insomnia (p<1e-4 and rg>0.5 for all traits), **Table 1 and Supplementary Table 6**). Finally, we found a strong association with different pain phenotypes, most notably back pain (rg=0.52, p=4e-5) and stomach and abdominal pain (rg=0.65,p=4e-5).

### Mendelian Randomization suggests that miserableness, nervous feelings are causal for nightmares

To explore causality between nightmares and psychiatric traits, we performed Mendelian Randomization with focus on phenotypes showing significant genetic correlation with the frequency of nightmares. Overall, psychiatric traits were also predictors for higher frequency of nightmares (feelings of fed up β=0.44 p=0.001; nervous feelings β=0.51, p=0.004 and miserableness β=0.53 p=0.0045) indicating that these traits contribute to nightmare frequency (Supplementary Table 7). No evidence of association or trend was seen that nightmares would be causal for any of the psychiatric traits (p=ns.).

## DISCUSSION

In this study we show that nightmares associate strongly with psychiatric and sleep traits at the genetic level. In addition, we show that psychiatric traits, notably those related to miserableness and nervous feelings can be causal for nightmares, while nightmares as such are not causal for the psychiatric traits. Furthermore, nightmares have a measurable genetic component and the genetic loci overlap with psychiatric and sleep complaints. In addition, individual genome-wide significant associations were discovered with *PTPRJ* and *MYOF*. Together with genetic correlations these findings strongly support the epidemiological findings that nightmares are an integral part of psychiatric traits and can even be caused by psychological problems.

An increased frequency of nightmares is seen among patients with psychiatric disorders ^8^. In the present study we found strong genetic correlations between nightmares and overall subjective well-being, mood disorders (major depression) and personality characteristics (neuroticism). Further, we identified genetic correlation between nightmares and sleep complaints; insomnia and excessive daytime sleepiness. This is in line with epidemiological findings that nightmares correlate with sleep and psychiatric traits ^3,4,8,29^. However, our findings suggest a novel mechanism where traits related to mood such as miserablesness and anxiety through nervous feelings are causal for nightmares. Notably, the current genetic correlations are computed in non-overlapping samples (Finnish population vs. UK Biobank) and therefore do not have shared individuals. Thus, the genetic correlation is not induced by overlapping phenotypic representation in the same individuals that would be part of the discovery sample.

One possible explanation for the co-occurrence of nightmares and psychiatric traits is the overlap between sleep complaints and nightmares, seen also in the present study. Nightmares occur usually during Rapid Eye Movement (REM) sleep, while instability of REM sleep accompanied by slower resolving of emotional distress has been found to be of key importance in the pathophysiology of primary insomnia ^30^. This could be one potential explanation for genetic and phenotypic correlations with nightmares. Our finding of shared genetic risk between nightmares and symptoms of insomnia would support this hypothesis.

We identified a novel individual risk variant for nightmares intronic for gene encoding *PTPRJ*. This locus has been previously associated with sleep duration and short sleep suggesting a possible larger effect of variation at the *PTPRJ* locus on sleep and nightmares. PTPRJ is a pleiotropic tyrosine phosphatase, previously implicated in regulation of angiogenesis, amount of platelets, cancer, blood glucose levels but also sleep duration and axonal growth^25,31-34^. In addition, two intronic variants for *MYOF*, the gene coding for *myoferlin*, associate with nightmares. The causal biological mechanisms for these variants remain still unknown and are potentially partially mediated through correlating psychiatric and sleep associations. Furthermore, the variants that we discovered in this study have remarkably different allele frequencies across populations making it challenging to assess causality and replication over multiethnic populations that will be affected by difference in exposures, population structure, selection or drift. Indeed, we will need larger samples with information on nightmares to replicate the findings related these genes, and to identify the causal variants, genes and causal mechanisms behind these genetic associations.

A subset of our findings may reflect that some individuals have a higher opportunity to recall dreams. For example, a higher nightmare recall frequency could be explained by higher number of awakenings during the night. Although the definition of a nightmare includes the awakening criterion, i.e., it is assumed that the dysphoric dream wakes up the dreamer, most individuals interpret nightmares simply as highly negative dreams and disregard the criterion for waking up. Also, in questionnaires, nightmares and the awakening criterion are seldom clearly defined for the participant. A higher dream and nightmare recall frequency could be explained by higher number of awakenings during the night, as this gives a greater opportunity to remember a dream on awakening, including nightmares. Some of our observations support this reasoning. Accordingly, we saw increased nightmares in individuals reporting night-time awakenings. On one hand, there was no genetic correlation with the number of sleep episodes. On the other hand, the strongest associating tissue enrichment in addition to brain was bladder suggesting that some individuals may need to get up during the night to use the bathroom, which also gives them an additional opportunity to recall dreams and nightmares.

Our study is unique as there are no other large samples available with phenotyping for nightmares, and is the first to examine genetic determinants of nightmares. Given the strong correlations with psychiatric and psychological traits it would be essential to grow awareness of nightmares affecting health and disease, systematically collect information about nightmares especially in clinical samples and implement these questions in larger cohorts. Our findings indicate a possibility that patients with psychiatric disorders may benefit if their nightmares are managed as part of the treatment strategy by using evidence-based interventions such as imagery rehearsal therapy^35^.

To summarize, these findings show two possible mechanisms for association of the genetic risk for nightmare with sleep traits (notably insomnia), either a direct mechanism that is tied to sleep problems and the negativity of a dream, or a modulatory effect that associates with the frequency that the dream is remembered. These data show that we need to implement questions for dream recall, nightmare frequency and distress, and night time habits also for large population samples in order to understand the connection between dreaming and nightmares in detail.

## URLs and data availability

The summary statistics for these associations will be available at Sleep disorder knowledge portal http://sleepdisordergenetics.org/.

## ACKNOWLEDGEMENTS

This study has been supported by the Academy of Finland grants #309643 Ollila, #290039 Paunio, CSC. NIH R01DK107859 and MGH Research Scholar Award to Saxena; Department of Defence through a National Defense Science and Engineering Grant and Stanford Graduate Fellowship Nasa Sinnott-Armstrong, HC has been supported by Finska Läkaresällskapet. JK has been supported by the Academy of Finland (grants 265240, 263278, 308248, 312073). Support for genotyping in the Finnish Twin Cohort has been provided by ENGAGE – European Network for Genetic and Genomic Epidemiology, FP7-HEALTH-F4-2007, grant agreement number 201413 and Wellcome Trust Sanger Institute.

## TABLES

Table 1. Genetic correlation between nightmares and strongest correlates from psychiatric and sleep disorders

Table 2. Individual GWAS hits associating with the frequency of nightmares

---

**Supplementary Figure 1.**
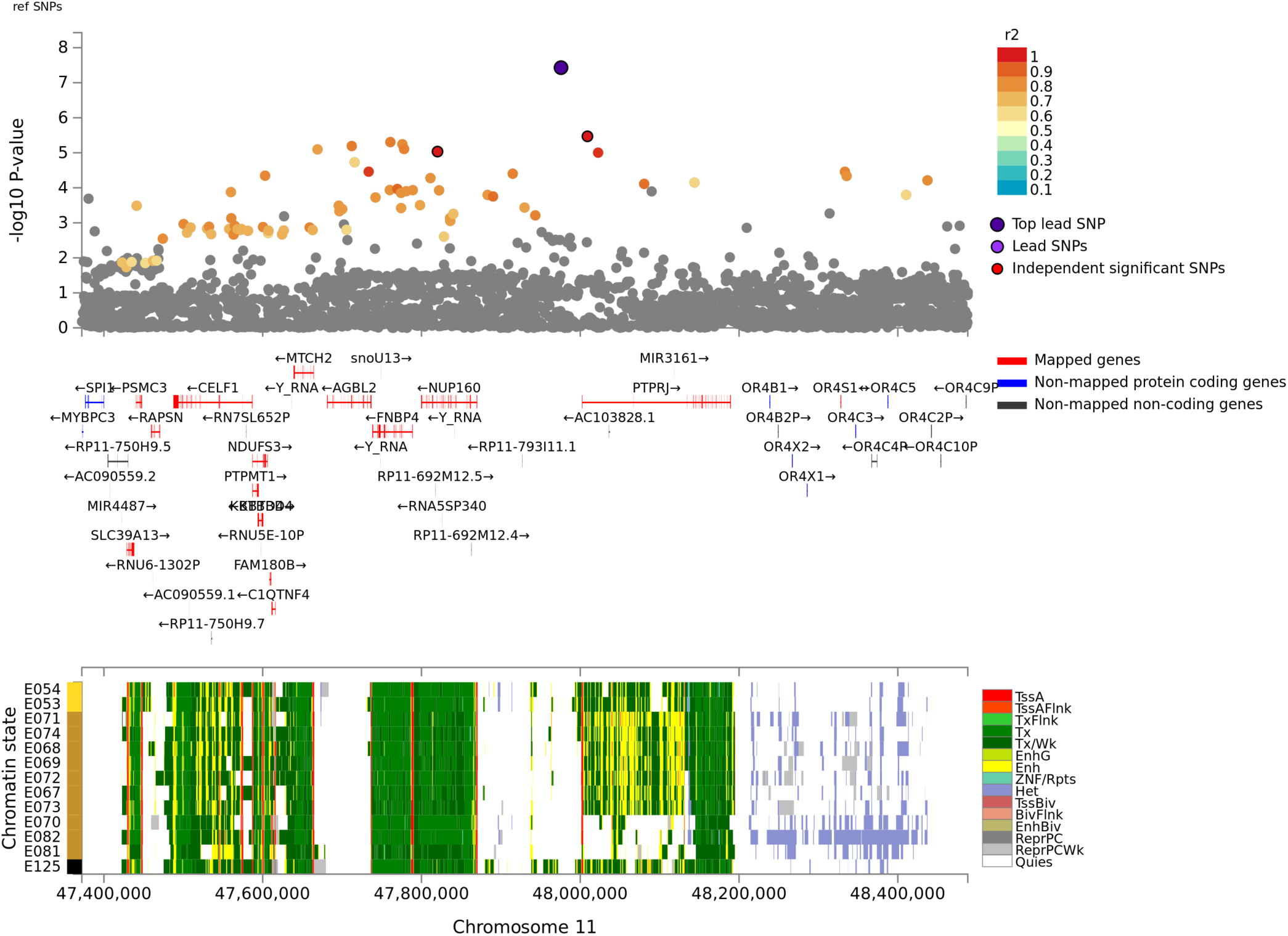
LocusZoom plot of *PTPRJ* locus and overlay with ENCODE 15 state model in Brain tissues.

**Supplementary Figure 2.**
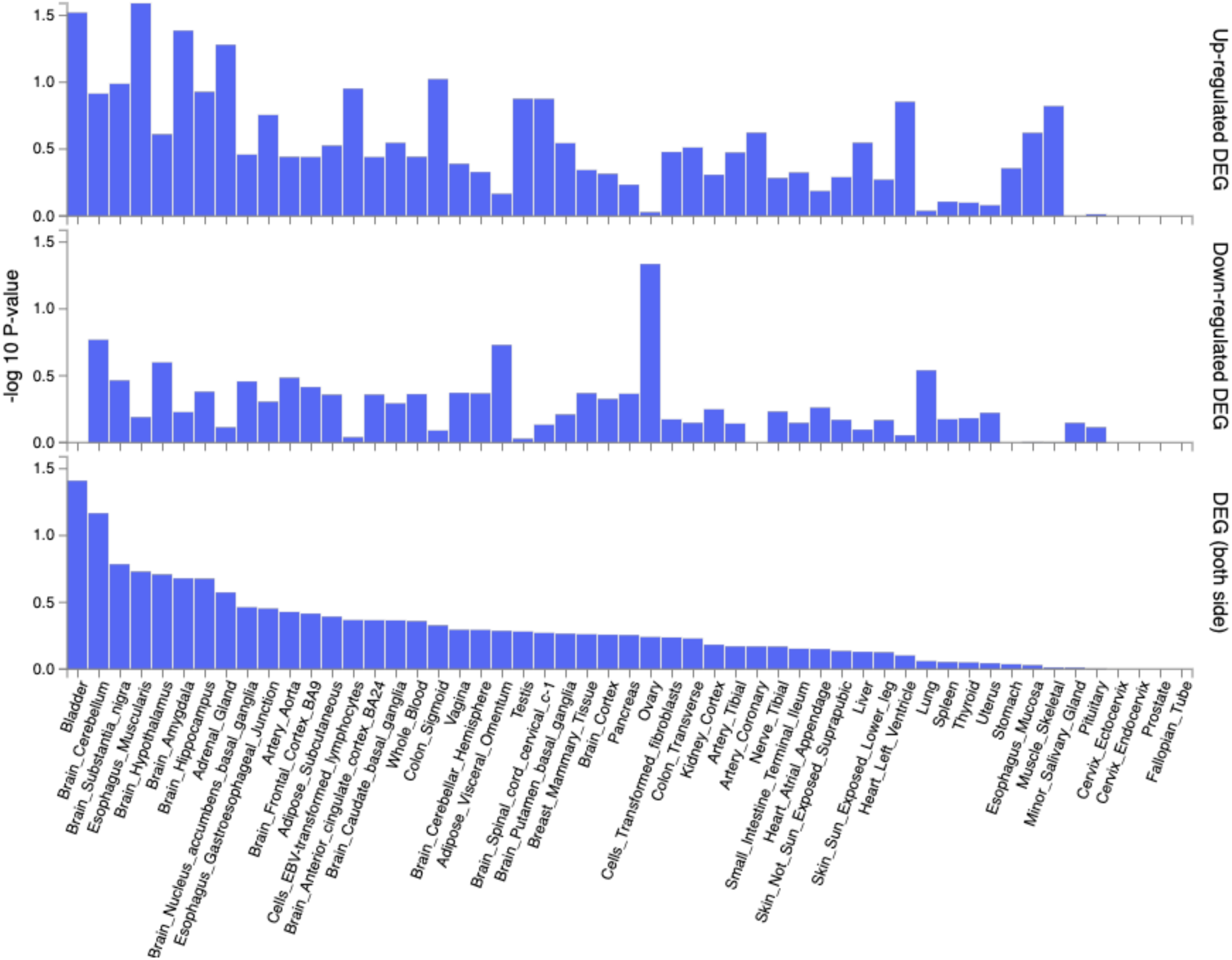
Enrichment of genes in tissues based on GTEx v6 expression.

